# Influenza virus-like particle-based hybrid vaccine containing RBD induces immunity against influenza and SARS-CoV-2 viruses

**DOI:** 10.1101/2022.02.01.478657

**Authors:** Ramireddy Bommireddy, Shannon Stone, Noopur Bhatnagar, Pratima Kumari, Luis E. Munoz, Judy Oh, Ki-Hye Kim, Jameson T. L. Berry, Kristen M. Jacobsen, Lahcen Jaafar, Swe-Htet Naing, Allison N. Blackerby, Tori Van der Gaag, Chloe N. Wright, Lilin Lai, Christopher D. Pack, Sampath Ramachandiran, Mehul S. Suthar, Sang-Moo Kang, Mukesh Kumar, Shaker J. C. Reddy, Periasamy Selvaraj

## Abstract

Several approaches have produced an effective vaccine against severe acute respiratory syndrome coronavirus 2 (SARS-CoV-2). However, the influence of immune responses induced by other vaccinations on the durability and efficacy of the immune response to SARS-CoV-2 vaccine is still unknown. We have developed a hybrid vaccine for SARS-CoV-2 and influenza viruses using influenza virus-like particles (VLP) incorporated by protein transfer with glycosylphosphatidylinositol (GPI)-anchored SARS-CoV-2 S1 RBD fused to GM-CSF as an adjuvant. GPI-RBD-GM-CSF fusion protein was expressed in CHO-S cells, purified and incorporated onto influenza VLPs to develop the hybrid vaccine. Our results show that the hybrid vaccine induced a strong antibody response and protected mice from both influenza virus and mouse-adapted SARS-CoV-2 challenges, with vaccinated mice having significantly lower lung viral titers compared to naive mice. These results suggest that the hybrid vaccine strategy is a promising approach for developing multivalent vaccines to prevent influenza A and SARS-CoV-2 infections.

## Introduction

Severe acute respiratory syndrome coronavirus 2 (SARS-CoV-2) first appeared in late 2019 in China before beginning its rapid spread across the globe (1). The disease, named COVID-19, presents a severe respiratory disease course and high fatality rate in the elderly and immunocompromised (2). The spike (S) protein of the virus binds to the human ACE2 protein for entry into epithelial cells of the respiratory tract (1, 3). This S protein, and specifically the conserved ACE2 receptor-binding domain (RBD), is a proven target for vaccine design. Antibodies and vaccines directed to the S protein and the RBD are effective in preventing SARS-CoV-2 infection (4). Virtually all recent vaccine strategies for SARS-CoV-2 target the full-length S protein (Pfizer/BioNTech, Moderna, Johnson & Johnson, AstraZeneca/Oxford, Novavax, Innovio, Curevac, and others) (5, 6).

The present study used a two-in-one hybrid vaccine for SARS-CoV-2 and influenza using a protein transfer method to incorporate a fusion protein of GM-CSF and RBD into influenza VLPs. The protein transfer approach achieves a higher level of incorporation of antigen and cytokine adjuvants in the VLP vaccine (7). Delivery of cytokine fusion protein antigens by the VLP to antigen presenting cells (APCs) enhances efficient presentation of the antigens to the immune system. Cytokines are known to increase the efficacy of vaccines by attracting and activating key immune cells (8–10).

Two cytokines under evaluation for their potential as biological adjuvants are granulocyte macrophage colony-stimulating factor (GM-CSF) and interleukin-12 (IL-12). In addition, targeting antigen to APCs via GM-CSF receptor enhances cross-priming (11, 12). GM-CSF potentiates a strong immune response primarily through maturation and differentiation of dendritic cells (13–16). The FDA approved a prostate cancer vaccine, Provenge®, by Dendreon, uses an antigen fused to GM-CSF that delivers antigen effectively to the immune system (17, 18). Adjuvants that have a tolerable safety profile and generate a Th1 immune response are of high importance. Moreover, proinflammatory cytokines produced by activated dendritic cells (DCs) have been shown to play an important role in inducing a robust immune response (19, 20). IL-12 has been well documented to induce a Th1 response, with a promising clinical benefit in cancer patients (8). IL-12, a heterodimeric cytokine (p35 and p40 subunits), activates DCs, T lymphocytes and natural killer (NK) cells to release IFN-γ, TNF-α etc. (21–23). IL-12 also induces T-cell precursors to differentiate towards a Th1 lineage, which also promotes development of a robust CTL response (24). Preclinical and clinical trials performed to evaluate the potential of recombinant soluble IL-12 as an adjuvant in treating several cancers and viral hepatitis and influenza resulted in enhanced immune response (9, 25–30), but also resulted in unfavorable side effects and systemic toxicity (31, 32).

However, delivering IL-12 in a membrane anchored form has been proved a successful approach in achieving desired adjuvant effect with minimal toxicity (33, 34). We engineered membrane-bound form of cytokines by attaching a GPI-anchor (35). The GPI-anchor permits incorporation of purified GPI-anchored proteins into the lipid bilayer of influenza VLPs or any amphiphilic micro/nano particles by a simple protein transfer technique (7, 36). By introducing the membrane incorporated GPI-cytokines into VLPs, multiple viral-specific antigens can be presented to the immune system to mount a robust immune response. In addition, administration of VLP vaccines containing membrane-anchored cytokines will localize the cytokines to the area of injection, thereby reducing the systemic effects associated with soluble cytokines (7). The VLP vaccine prepared by our protein transfer approach requires only a low amount of GPI-GM-CSF (0.025 μg/μg of VLP) for optimum antiviral response in mice. Moreover, physical linkage of adjuvant and antigen source leads to simultaneous adjuvant and antigen delivery to immune cells, resulting in enhanced immune reactivity and increased vaccine efficacy when compared to unconjugated antigen and adjuvant mixture (37, 38). In the present study, we demonstrate that influenza VLP vaccine incorporated with GPI-RBD-GM-CSF fusion protein and GPI-IL-12 (hybrid vaccine) protected mice from influenza virus challenge and also induced a robust, durable antibody response in BALB/c mice as well as decreased viral load and less weight loss when challenged with mouse adapted SARS-CoV-2.

## RESULTS

### Characterization of GPI-RBD-GM-CSF fusion protein

We constructed a fusion protein gene by joining the DNA sequences specific to SARS CoV-2 S protein RBD domain, mouse GM-CSF sequence and the GPI-anchor signal from human CD59 (Figure 1A) and expressed the gene in CHO-S cells. Flow cytometry analysis demonstrated expression of the RBD-GM-CSF fusion protein on the surface of transfected CHO-S cells (Figure 1B). To verify whether the GPI-RBD-GM-CSF fusion protein contains the GPI tail, CHO-S cells expressing the fusion protein were incubated with phosphatidylinositol-specific phospholipase C (PIPLC), which is known to cleave the GPI moiety from the proteins and release them from the cell membranes, and then analyzed by flow cytometry. The results show that nearly 76% of the fusion protein was released from the cell surface, suggesting that the fusion protein is anchored to the cell membrane via GPI tail (Figure 1C). To determine whether the fusion protein on the CHO-S cell surface binds to its cognate receptor ACE2, flow cytometry analysis was used to detect the binding of biotinylated human ACE2 (ACRO Biosystems) to the CHO-S cells. The results verified that the RBD in the fusion protein retained its ACE-2 binding activity (Figure 1D). Neutralizing antibody MM57 (Sino Biologicals) was able to bind to GPI-RBD-GM-CSF on CHO-S cell transfectants, which was blocked by pre-incubation with purified GPI-RBD-GM-CSF and commercially available Spike RBD (RayBiotech), confirming that the fusion protein retains the RBD conformation (Fig. S1). Transfected cells were cloned and CHO-S clone 3C3 was used for further studies.

**Figure 1.**
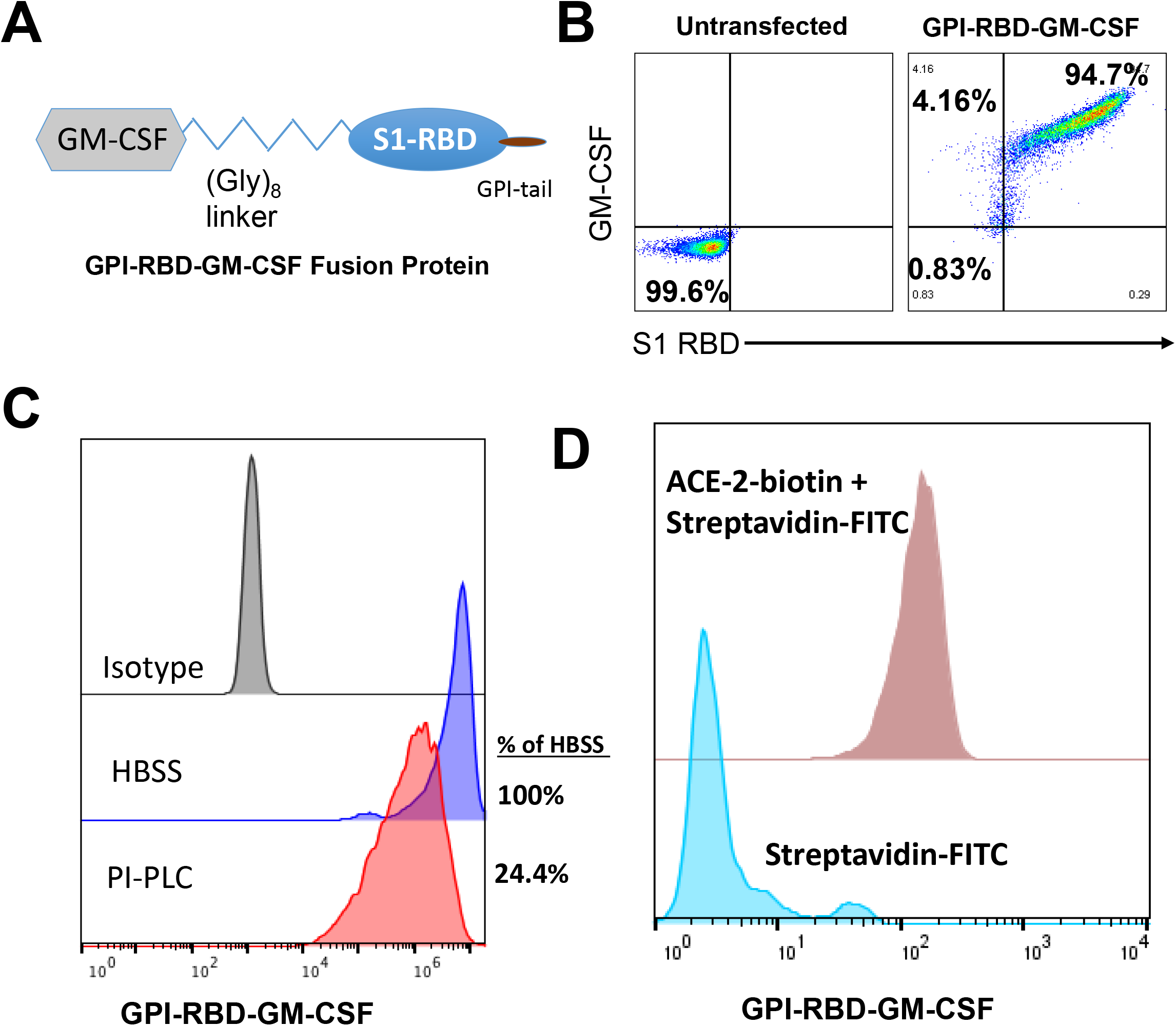
Design, expression, and characterization of GPI-RBD-GM-CSF fusion protein. (**A**) Design of GPI-RBD-GM-CSF fusion protein gene, (**B**) GPI-RBD-GM-CSF fusion protein binds both anti-RBD mAb and anti-GM-CSF mAb on CHO-S cell transfectants, (**C**) PIPLC treatment of CHO-S cells expressing GPI-RBD-GM-CSF reduced the level of expression, and (**D**) Flow cytometry analysis showed binding of human ACE-2 to GPI-RBD-GM-CSF fusion protein expressed in CHO-S cells.

### GPI-RBD-GM-CSF fusion protein retains functional activity

CHO-S cell expressed GPI-RBD-GM-CSF fusion protein was affinity purified using NHS-Sepharose coupled to anti-mouse GM-CSF antibody. The purified fusion protein was run on 12 % SDS-PAGE under non-reducing conditions and either stained with Colloidal blue (Figure 2A, lane 2) or detected with western blots using anti-RBD antibody (Figure 2A, lanes 3 & 4), and anti-GM-CSF (Figure 2A, lanes 6 & 7). Affinity purified GPI-RBD-GM-CSF fusion protein runs as a smear ranging from 50 kDa (size of non-glycosylated fusion protein) to 250 kDa with several distinct bands with sizes 55 kDa, 110 kDa and 220 kDa. Western blot analysis was performed for identity and size comparison of the fusion protein to wild-type mouse GPI-GM-CSF (Figure 2A, lane 5). Most of the colloidal blue stained bands were detected by anti-mouse GM-CSF and anti-RBD antibodies suggesting that they are multimeric forms of the fusion protein.

**Figure 2.**
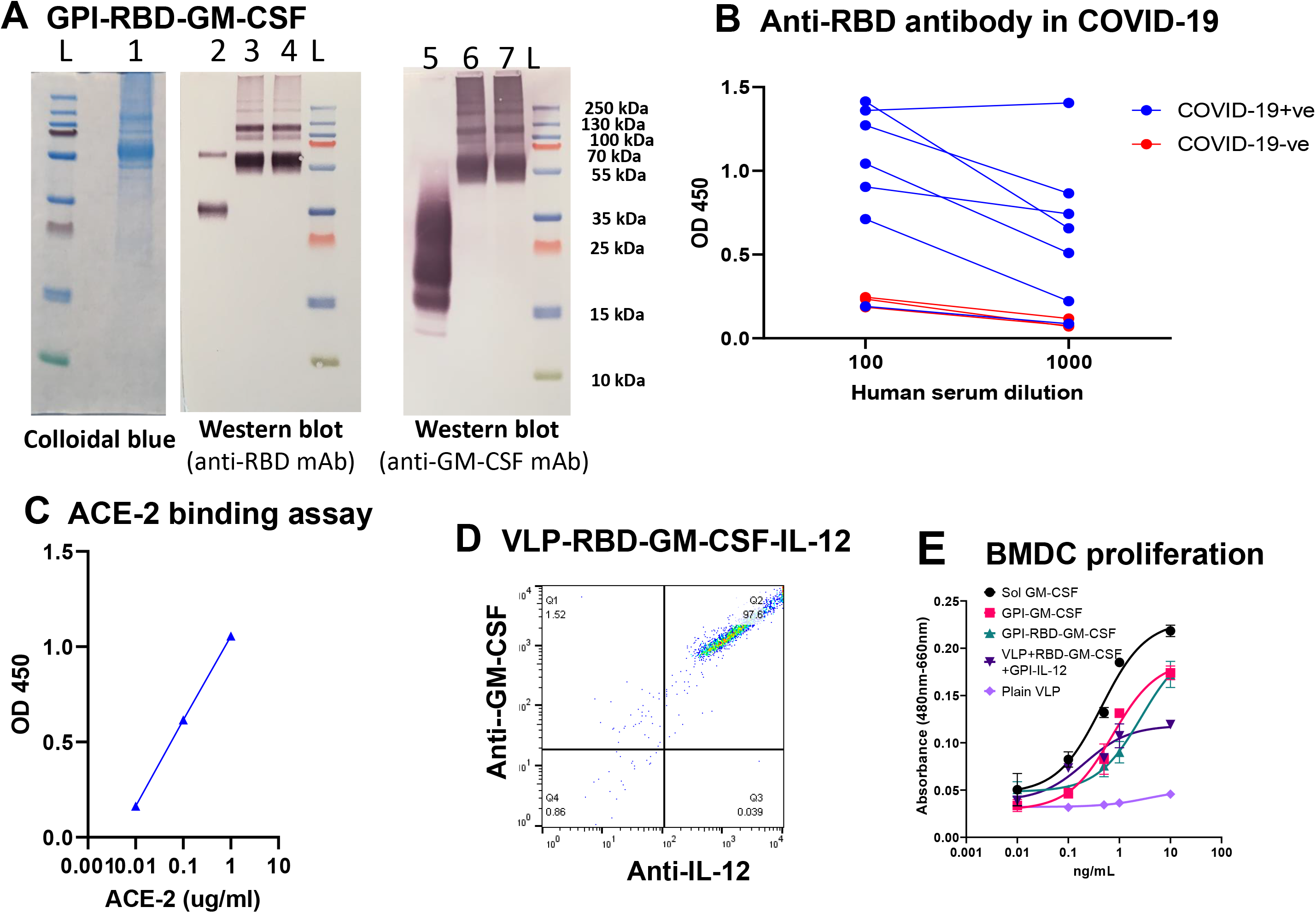
Purified GPI-RBD-GM-CSF fusion protein retains functional activity. (**A**) Colloidal blue (lane 1), and Western blot (lanes 2-7) of the immunoaffinity column purified fusion protein from CHO-S cells probed with anti-RBD antibody (lanes 3 & 4), or anti-GM-CSF mAb (lanes 6 & 7). Lane 2 is control RBD probed with anti-RBD Ab, and lane 5 is control GM-CSF probed with anti-GM-CSF antibody. (**B**) ELISA for GPI-RBD-GM-CSF binding to antibodies in the human COVID-19 patients’ sera, and (**C)** ELISA of purified GPI-RBD-GM-CSF binding to ACE-2. (**D**) FACS analysis of VLPs incorporated with GPI-IL-12 and GPI-RBD-GM-CSF fusion protein by protein transfer, and (**E**) BMDC proliferation by GPI-GM-CSF, GPI-RBD-GM-CSF and hybrid vaccine.

To test whether the anti-RBD antibodies in the convalescent sera from human patients (RayBiotech) recognize the RBD in the purified GM-CSF fusion protein, we performed a direct ELISA. The data show that antibodies from COVID-19 positive sera that bind spike protein-RBD bind to the GPI-RBD-GM-CSF (Figure 2B). Next, we tested whether the purified fusion protein has dual function as GM-CSF and also binds human ACE2. An ELISA using biotinylated human ACE2 confirmed that the RBD in purified GPI-RBD-GM-CSF fusion protein retains its receptor binding activity (Figure 2C).

To make the VLP hybrid vaccine, we incorporated the influenza VLPs with GPI-IL-12 and GPI-RBD-GM-CSF by protein transfer. Influenza VLPs contain a lipid bilayer which is amenable to protein transfer mediated incorporation of GPI-proteins. Flow cytometry was performed using fluorochrome-conjugated anti-IL-12 and anti-GM-CSF antibodies for confirming the dual incorporation of the GPI-RBD-GM-CSF and GPI-IL-12 onto VLPs (Figure 2D). Western blot analysis of the hybrid vaccine using anti-IL-12, anti-GM-CSF and anti-RBD antibodies confirmed the presence of GPI-IL-12 and GPI-RBD-GM-CSF fusion protein in the hybrid vaccine (data not shown). To test whether VLP-incorporated GM-CSF retains its function, we cultured mouse bone marrow derived dendritic cells (BMDC) *in vitro* with soluble GPI-RBD-GM-CSF or VLPs incorporated with GPI-RBD-GM-CSF and measured BMDC proliferation using XTT assay. The results show that GPI-RBD-GM-CSF incorporated in the VLP vaccine is capable of stimulating BMDC proliferation (Figure 2E).

### Hybrid vaccine induces durable antibody response

To test whether the GPI-RBD-GM-CSF fusion protein induces antibody response, we immunized BALB/c (2-3 months old) mice with GPI-RBD-GM-CSF fusion protein (0.1, 1.0, 2.0 and 5.0 μg) or VLPs (1.0, 2.0, 5.0 and 10 μg) incorporated with the fusion protein and GPI-IL-12. Controls included VLP without cytokines, commercially available RBD (RBD-His from Ray Biotech) or PBS. Booster dose was given 2-4 weeks after the first dose. The route of administration was either subcutaneous (s.c.) or intramuscular (i.m.) (Fig. S2).

Blood was collected every 2 to 4 weeks for antibody titer, ACE2 binding inhibition, and virus neutralization assays. VLP vaccine incorporated with GPI-RBD-GM-CSF and GPI-IL-12 (hybrid vaccine) induced robust antibody response after the booster dose (Fig. S3). The antibody response against GPI-RBD-GM-CSF fusion protein was comparable in both i.m. and s.c. routes of vaccine administration (Fig. S4). The antibodies bind to CHO-S cells expressing the GPI-RBD-GM-CSF fusion protein but not untransfected CHO-S cells (Fig. S5) confirming the specificity of the antibody response to GPI-RBD-GM-CSF fusion protein. The antibody response was durable at least for 6 months against RBD (Fig. S6A), and hemagglutination inhibiting antibodies against A/PR8 influenza virus were even longer, present up to 15 months after booster dose (Fig. S6B and C), showing that VLP-based vaccine is effective in inducing durable RBD specific and influenza virus specific HAI antibody responses.

### Hybrid vaccine induces SARS-CoV-2 neutralizing antibodies

To test whether antibodies generated by the hybrid vaccine block the binding of Spike RBD to human ACE-2, we performed an inhibition assay using biotinylated ACE-2 and CHO-S cells expressing GPI-RBD-GM-CSF fusion protein. Our results show that antibodies induced by both purified fusion protein and hybrid vaccine blocked ACE2 binding to RBD on CHO-S cells (Fig. S7). We have used ACE-2 blocking anti-RBD (clone MM57) mAb (2 μg and 10 μg) as a positive control. For live virus neutralization, Vero6 cells and WA1 strain of SARS-CoV-2 were used in a modified FRNT assay as described in a previously published study (39). While the purified GPI-RBD-GM-CSF induced levels of antibody response similar to that of VLP hybrid vaccine (Fig. S8A), neutralizing antibody titers were very low in the mice that received only the purified fusion protein GPI-RBD-GM-CSF without VLP. The data suggest that VLP incorporated fusion protein induced stronger neutralizing antibodies than the soluble fusion protein (Fig. S8B).

### Hybrid vaccine induces IgG2a antibody response

We observed that purified GPI-RBD-GM-CSF fusion protein by itself or incorporated onto VLPs induced a strong antibody response (Figure 3A). However, recombinant RBD-His tag failed to induce an antibody response suggesting that the GM-CSF in our fusion protein is acting as an adjuvant (Fig. S9). To determine the isotype of antibodies induced by the hybrid vaccine, we have performed antibody isotyping ELISA as described in the Methods section. While the purified GPI-RBD-GM-CSF fusion protein alone induced an elevated antibody response, mostly Th2 type IgG1 (blue circles, Figure 3B), the hybrid VLP vaccine induced both IgG2a (a Th1-induced response) and IgG1 (orange, red and green symbols, Figure 3B). However, addition of GPI-IL-12 to the VLP vaccine did not further enhance the IgG2a response (green versus red bars).

**Figure 3.**
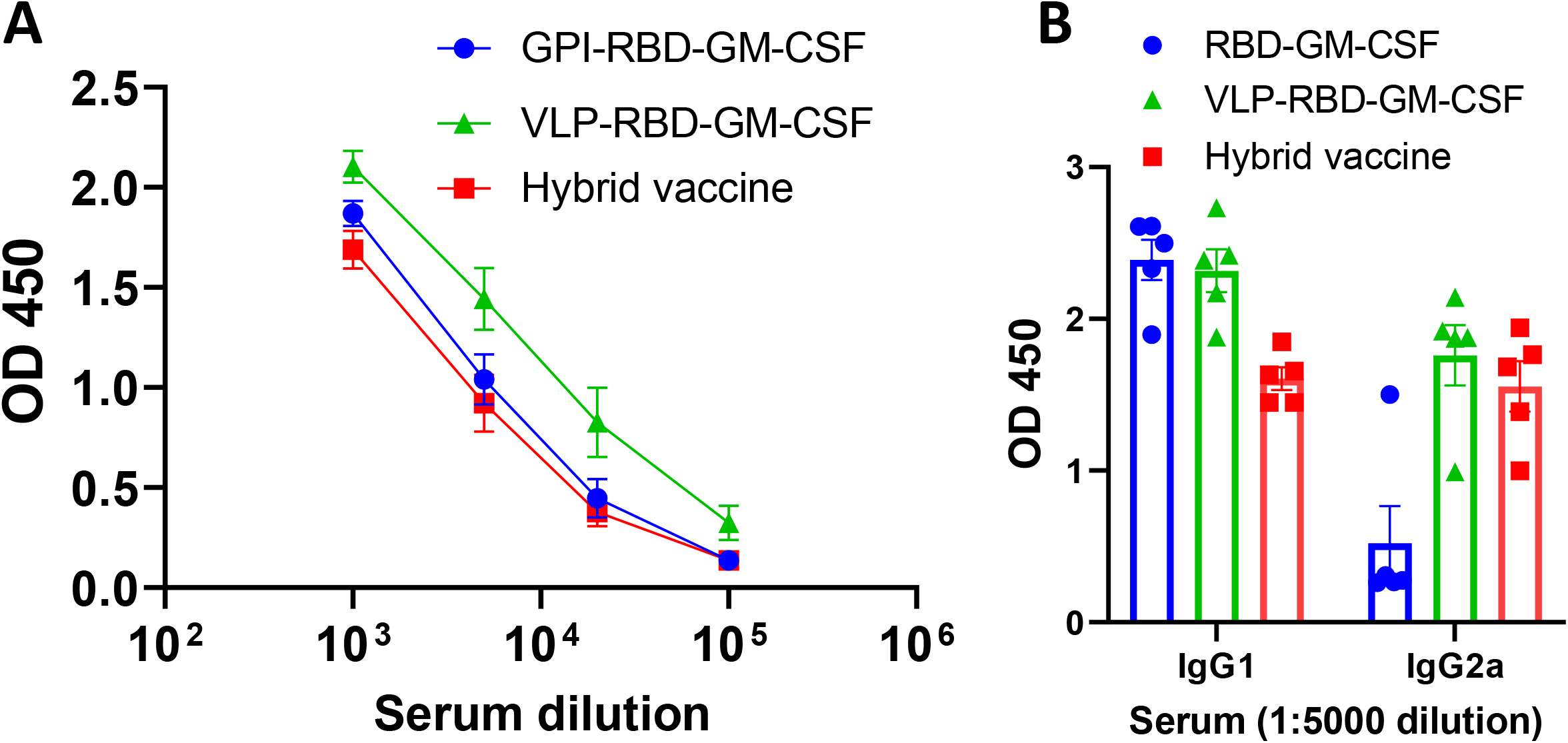
Hybrid vaccine induces antibody response against Spike RBD in mice. ELISA plates were coated with GPI-RBD-GM-CSF and serum samples from various groups of mice (n=5) immunized with GPI-RBD-GM-CSF, VLP vaccine containing the GPI-RBD-GM-CSF or hybrid vaccine (VLP vaccine containing the GPI-RBD-GM-CSF + GPI-IL-12) were diluted and added to the wells after blocking the plates. (A) Total IgG levels 6 weeks after booster dose, and (B) IgG isotype in the sera 10 weeks after the booster dose. Anti-mouse IgG (A) or isotype specific anti-mouse IgG-HRP conjugate (B) was used to detect the bound antibody.

### Hybrid vaccine induces anti-influenza virus IgG1 and IgG2a antibody response

To test whether the hybrid vaccine induced antibodies against influenza virus antigens, we analyzed the sera for antibodies and isotype of the antibodies. Our results show that VLP vaccine with GPI-RBD-GM-CSF with and without GPI-IL-12 induced equally potent antibody responses against inactivated influenza A/PR8 virus (data not shown). VLP vaccine incorporating the cytokines (purple and red symbols) induced higher levels of antibodies compared to VLP without cytokines (blue symbols, Figure 4A). The antibody response is a mixed Th1 and Th2 type (IgG1 and IgG2a) against influenza A/PR8 antigens in inactivated virus (Figure 4A - C) and influenza A/PR8 VLP (Fig.S10). Interestingly, we observed that VLP vaccine without RBD-GM-CSF that was incorporated with GPI-GM-CSF instead (purple symbols) induced significantly higher levels of IgG2a (Figure 4C) but not IgG1 (Figure 4B) compared to the VLP administered group suggesting that GPI-IL-12 and GPI-GM-CSF in the VLP vaccine are augmenting Th1 type IgG2a antibody response. The incorporation of GPI-RBD-GM-CSF instead of GPI-GM-CSF onto the VLP vaccine diminished the level of anti-influenza antibody response (red symbols in Figure 4C). This may be because GPI-GM-CSF is more active than GPI-RBD-GM-CSF or may be due to the difference in the level of incorporation onto VLP or both, which need further investigation.

**Figure 4.**
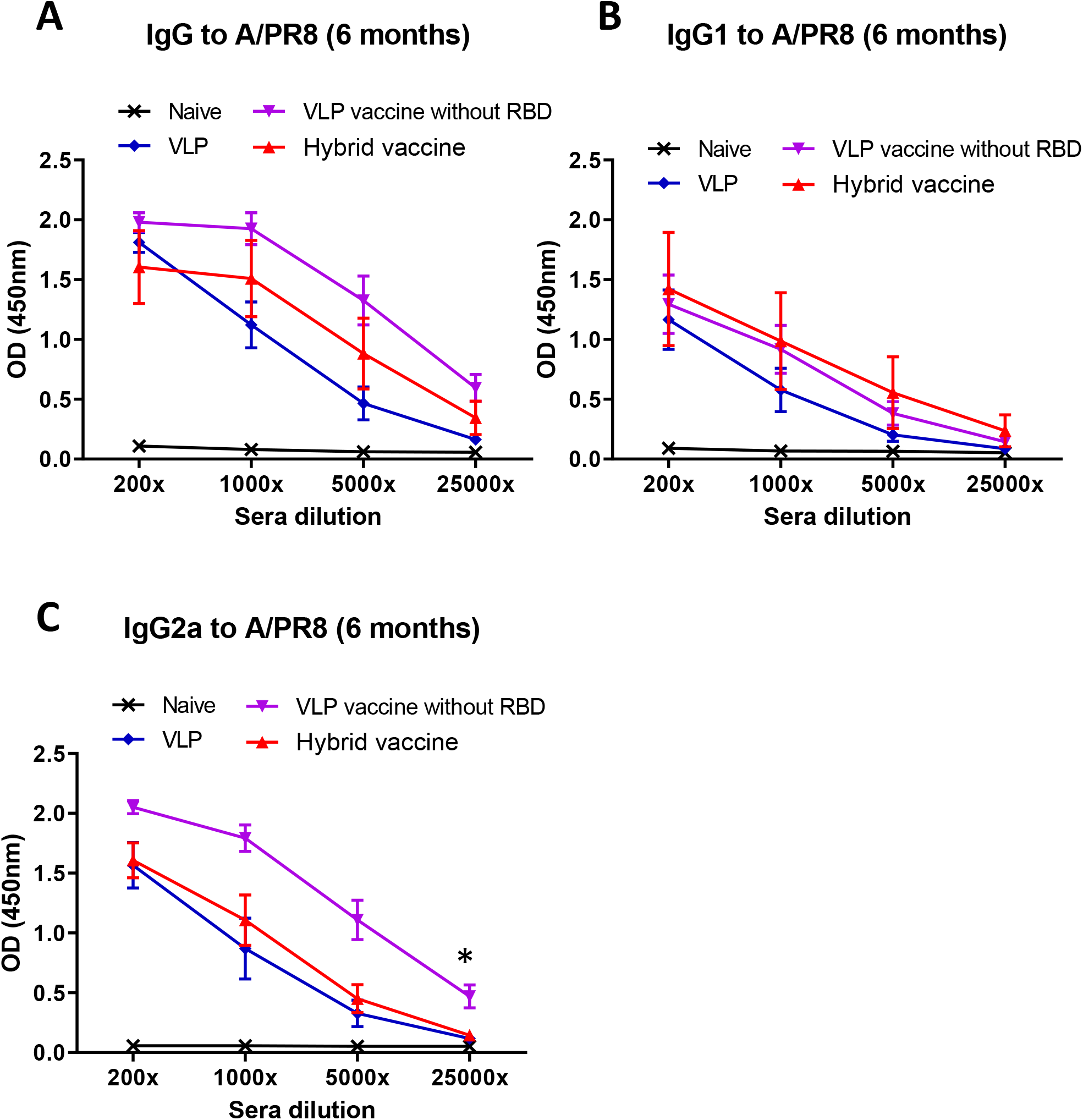
Hybrid vaccine induces influenza A/PR8 virus-binding IgG1 and IgG2a antibody response. ELISA plates were coated with inactivated influenza A/PR8 H1N1. Serum samples from mice vaccinated with VLP, VLP with GPI-GM-CSF and GPI-IL-12 or hybrid vaccine were serially diluted (5-fold) and added to the wells after blocking the plates. (**A**) total IgG, (**B**) IgG1 or (**C**) IgG2a antibody response. Isotype specific anti-mouse Ig-HRP conjugate was used to detect the isotype of the antibody bound. * p<0.05

### Hybrid vaccine protects mice from SARS-CoV-2 and influenza virus challenges

To test whether the vaccine administration induces protective response, mice were immunized with unmodified VLP, hybrid vaccine without IL-12 (VLP RG) and hybrid vaccine. A booster dose was given after 33 days of first dose. Blood was collected 2 weeks after the booster dose and anti-RBD antibody response against RBD and influenza virus antigens, and neutralizing antibody titers using live SARS-CoV-2 (Wuhan strain) were performed as described in Methods section. Our results indicate that the hybrid vaccine and VLP RG induced strong antibody responses against RBD and influenza VLP antigens (Fig. S11), and the antibodies were able to neutralize live virus infection in a plaque reduction neutralization titer (PRNT) assay (Figure 5A). These mice were transferred to a ABSL3 facility for challenging with mouse adapted SARS-CoV-2 virus (MA10) after confirming the neutralizing antibody titers. Mice were challenged with 10^5^ plaque forming units of the SARS-CoV-2 virus. We observed acute weight loss in control VLP administered mice but the mice that were administered with either hybrid vaccine or hybrid vaccine without IL-12 (VLP RG) were protected from weight loss (Figure 5B). Since this virus is not lethal, mice were euthanized for quantification of lung viral titer after 3 days of challenge. Virus titer estimates revealed that hybrid vaccine or VLP RG decreased the virus replication significantly compared to the mice that received VLP. Hybrid vaccine that contained IL-12 in addition to RBD-GM-CSF fusion protein was more effective in controlling lung viral loads than VLP-RBD-GM-CSF (VLP RG) that lacked IL-12 (Figure 5C).

**Figure 5.**
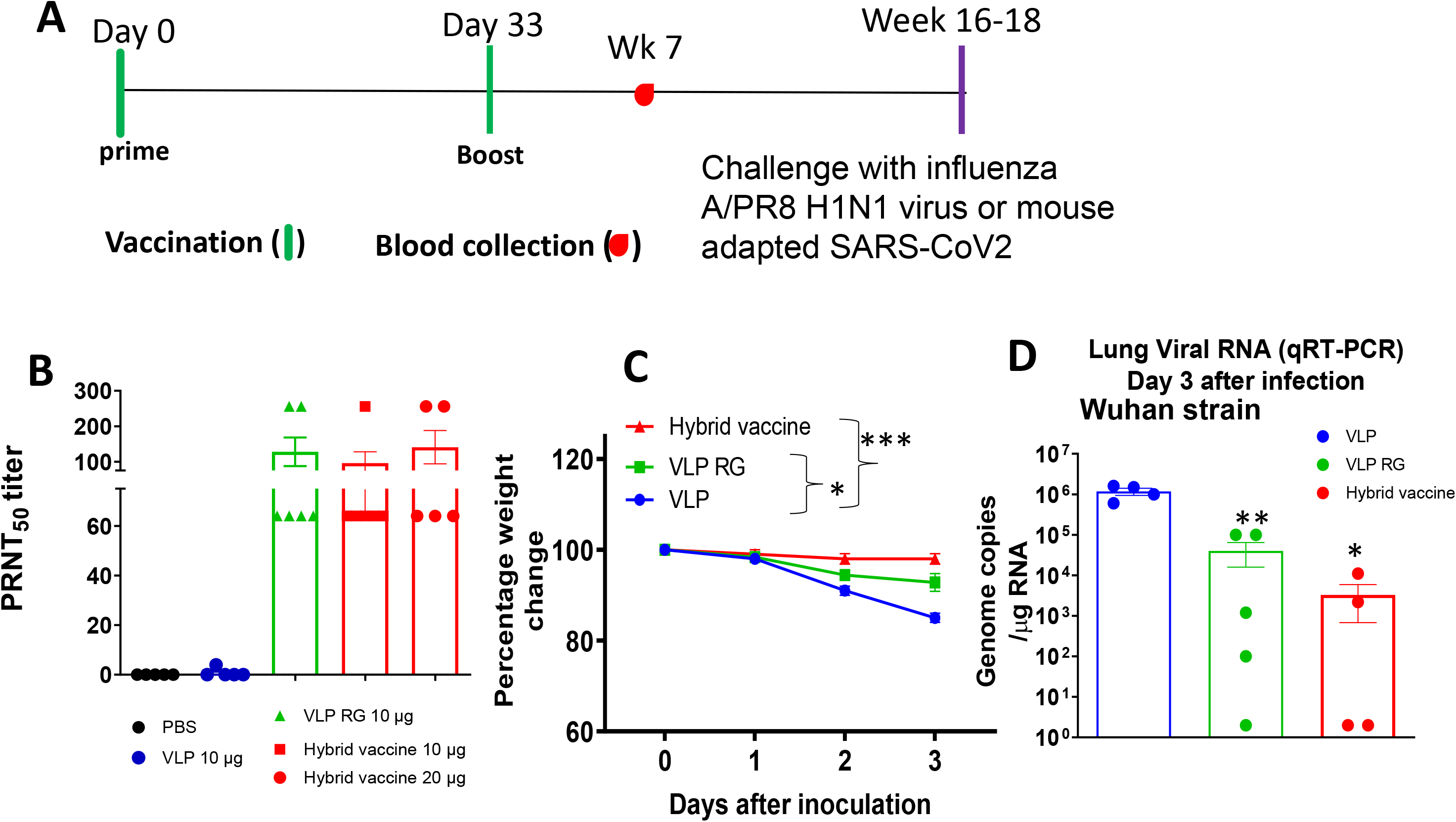
Hybrid vaccine protects against SARS-CoV-2 challenge. BALB/c mice (n=10; 2-3 months old) administered with hybrid vaccine, hybrid vaccine without IL-12 (VLP RG) or VLP in 50 ul volume via intramuscular route. Control mice received either PBS or influenza VLP. Booster dose given on d33, and blood was collected 2 weeks later (week 7). Mice were challenged with influenza A/PR8 H1N1 virus (n=5) or mouse adapted SARS-CoV2 (n=5) 16-18 weeks after the first dose. (**A**) Neutralizing antibodies titers in the serum of BALB/c mice (n=5-6 per group). Serum collected from BALB/c mice 2 weeks after booster dose was serially diluted from 1:4 to 1:1024 and PRNT was conducted against SARS-CoV-2 (Wuhan virus). (**B**) BALB/c mice were inoculated intranasally with mouse-adapted SARS-CoV-2 (10^5^ plaque-forming units) 3 months after booster dose. Percentage of daily body weight change in the animals. (**C**) The RNA levels of SARS-CoV-2 were determined in the lungs by qRT-PCR (n=4-5 per group). Error bars represent SEM. The data are expressed as genome copies/μg of RNA. Each data point represents an individual mouse. Data are expressed as mean log_10_ titer. * p<0.05; ** p <0.01 ***p<0.001. VLP RG: VLP incorporated with GPI-RBD-GM-CSF; hybrid vaccine: VLP incorporated with GPI-RBD-GM-CSF and GPI-IL-12.

Another cohort of mice from the same treatment groups were transferred to ABSL2 facility for influenza A/PR8 virus challenge. Hemagglutination inhibition titers against A/PR8 virus were determined before challenge. The results showed that VLP, VLP RG and hybrid vaccine administration induced high titers of hemagglutination inhibiting antibodies (Figure 6A). Mice were challenged with 10 LD_50_ of influenza A/PR8 and monitored for their body weight changes and survival rates. Since the mice administered with VLP or VLP vaccines were protected from weight loss compared to control infected mice (data not shown), we have euthanized 3 mice from each group after 5 days of infection and measured A/PR8 H1N1 titers in the lungs. We observed that the viral loads were significantly reduced up to a level of detection limit in mice that received VLP, hybrid vaccine or VLP RG (Figure 6B) compared to the PBS control. The virus was undetectable in mice administered with VLP RG or hybrid vaccine, but VLP alone was also equally protective against influenza virus. Consistent with the viral titers, the mice vaccinated with VLP or VLP with cytokine adjuvants survived lethal challenge (Figure 6C).

**Figure 6.**
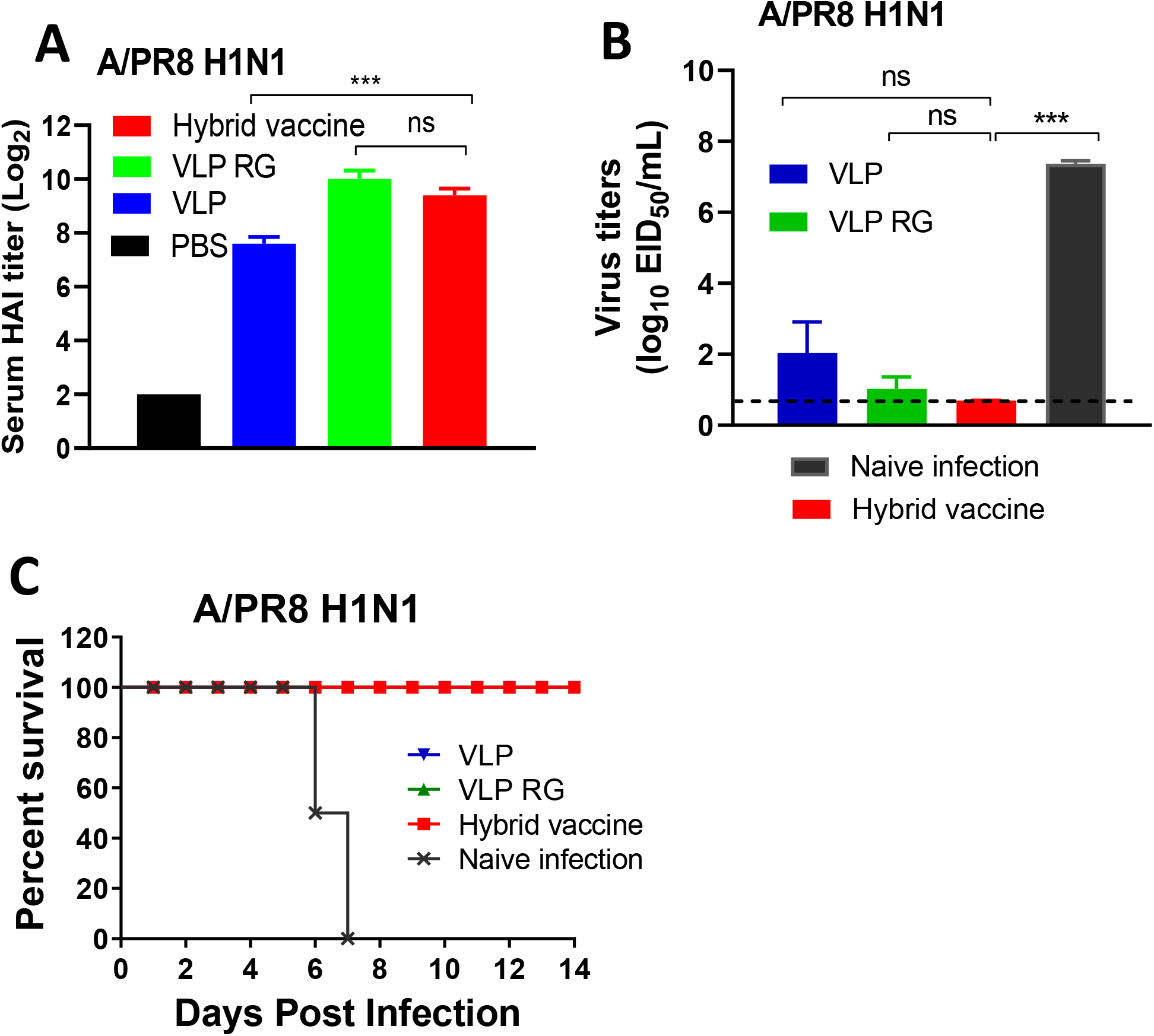
Hybrid vaccine protects against influenza A virus challenge. BALB/c mice administered with hybrid vaccine (10 μg/dose) or hybrid vaccine without GPI-IL-12 (VLP RG) as described in Figure 5. Booster dose given after 33 days of the first dose. (**A**) HAI titer in the blood 2 weeks after the booster dose (day 48). Lung viral titer (**B**) and survival (**C**) of mice challenged with influenza A/PR8 H1N1 virus 3 months after the booster dose. Inoculation dose was 10 times LD50. VLP RG: VLP incorporated with GPI-RBD-GM-CSF; Hybrid vaccine: VLP incorporated with GPI-RBD-GM-CSF and GPI-IL-12. Statistical significance was calculated by one-way ANOVA and Dunnett’s post-multiple comparison tests. Error bars indicate the mean ± standard errors of the mean (SEM). ***; p < 0.001, ns; not significant

## DISCUSSION

We have demonstrated that a hybrid vaccine based on influenza VLPs induces effective immunity against SARS-CoV-2 and influenza viruses. Our vaccine platform is based on cytokine adjuvants linked to the VLPs that carry the antigens (7). This approach delivers both antigens and biological adjuvants to the immune system simultaneously in a particulate form. We have generated CHO-S cells expressing GPI-anchored spike RBD-GM-CSF fusion protein and GPI-IL-12, purified the proteins and incorporated them onto influenza VLPs to develop the hybrid vaccine. Administration of the hybrid vaccine via either subcutaneous or intramuscular routes induced comparable levels of antibody response. Since soluble GPI-RBD-GM-CSF induced antibody response, but not the recombinant spike RBD without GM-CSF, our results suggest that GM-CSF in the fusion protein is acting as an adjuvant. Our approach to use cytokine adjuvants targets the antigen to APCs via their receptors and also allows the cytokines to enhance APC maturation. GM-CSF alone acting as an adjuvant in our GPI-RBD-GM-CSF fusion protein can induce a robust antibody response. Interestingly, the antibody response induced by the purified fusion protein is primarily the non-neutralizing IgG1 isotype, whereas fusion protein delivered using VLPs induced a neutralizing IgG2a and IgG1 mixed isotype antibody response. Our results are consistent with the recent report by the Bjorkman laboratory (40) and our previous studies on tumor antigens (36) demonstrating the use of VLP as a delivery vehicle for antigens for inducing a protective immune response. The hybrid vaccine also induced neutralizing antibodies against influenza A/PR8 (H1N1), suggesting that this approach of delivering spike RBD on an influenza VLP along with cytokine adjuvants confers dual protection against both influenza A H1N1 and SARS-CoV-2 viruses.

Mice challenged with H1N1 live virus 3 months after the booster dose were still well protected, suggesting that anti-influenza antibody and T cell responses induced by hybrid vaccine are long lasting. Neutralizing antibody titers against inactivated influenza A/PR8 (H1N1) are high even after 6 months of vaccination, also confirming the durability of the anti-influenza immune response induced by hybrid vaccine. Mouse adapted SARS-CoV-2 virus infection causes body weight loss but does not cause lethality in BALB/c mice (41). This finding was observed within 3 days of infection in control mice that were vaccinated with VLP, and hybrid vaccine prevented mice from losing weight. Lung virus titers were significantly decreased in mice that were vaccinated with hybrid vaccine compared to plain VLP. This suggests that the hybrid vaccine containing GPI-RBD-GM-CSF with cytokine adjuvants confers protection from severe disease caused by SARS-CoV-2 infection. Administration of purified GPI-RBD-GM-CSF fusion protein without VLP also induced long lasting antibody response, observed even after 1 year of booster dose. However, the antibodies were not able to neutralize live virus even though they were able to block ACE-2 binding to Spike S1. The antibodies are mostly IgG1 in mice vaccinated with purified GPI-RBD-GM-CSF fusion protein alone whereas hybrid vaccine (VLP with GPI-RBD-GM-CSF fusion protein and GPI-IL-12) induced both IgG1 and IgG2a isotypes, and blocked SARS-CoV-2 virus infection. This suggests that Th1 type response induced by the hybrid vaccine is more protective than Th2 type response induced by the purified GPI-RBD-GM-CSF.

The protein transfer approach used here to prepare hybrid vaccine allows anchoring of these cytokines to the surface of the VLPs which limits the systemic toxicity of the cytokines by acting as depot at the site of vaccination. Physical linkage of adjuvant and antigen source results in presentation of the adjuvant and antigen simultaneously to the immune cells, leading to enhanced immune reactivity and increased vaccine efficacy. Such a physical linkage of antigen and adjuvants is more effective than unconjugated antigen and adjuvant mixture (37). In addition, IL-12 and GM-CSF target antigen presenting cells, such as dendritic cells, by binding to IL-12 and GM-CSF receptors and enhancing antigen uptake and presentation, thereby enhancing subsequent T cell responses.

The limitation of the current study is that a comparative analysis of GPI-RBD-GM-CSF and GPI-RBD was not carried out to demonstrate the contribution of GM-CSF as an adjuvant in VLP vaccine. Attempts to make GPI-RBD in CHO cells were not successful. Further, our study did not investigate whether incorporating VLPs with different amounts of GPI-anchored fusion protein results in stronger immune response to SARS-CoV-2. However, the comparison of antibody response induced by purified GPI-RBD-GM-CSF molecule with RBD-His-Tag suggests that GM-CSF stimulated antibody production against RBD. Further, our study focused only on the RBD domain of SARS-CoV-2 S protein but not the full-length S protein which may limit the breadth of protective immune response.

In summary, our results demonstrate that influenza VLP-based delivery of SARS-CoV-2 RBD protein in combination with cytokine adjuvants can be used as a platform to develop multivalent vaccines targeting the variant strains of viruses which are currently observed in ongoing SARS-CoV-2 pandemic. Our fusion protein vaccine design also allows for creation of fusion proteins with new variant sequences and quickly purify using anti-GM-CSF mAb affinity chromatography. Further, use of immobilized cytokines as adjuvants will provide a safer way to induce anti-viral immunity with minimal side-effects.

## MATERIALS AND METHODS

### Antibodies and proteins

Purified anti-mouse GM-CSF (clone MP1-22E9) and anti-mouse IL-12 (clone C17.8) were from BioXcell and used for affinity chromatography purification of GPI-RBD-GM-CSF fusion protein and GPI-IL-12, respectively. Anti-spike RBD antibody (clone MM57) obtained from Sino Biologicals (Cat#40592). FITC-conjugated goat secondary antibody against mouse IgG/IgM was purchased from BD Pharmingen (Cat# 555988). Peroxidase (HRP)-conjugated goat anti-mouse IgG F(ab’)_2_ specific antibody was from ThermoFisher Scientific/Pierce (Cat#31436). Antibody isotyping kit was purchased from Southern Biotech (Cat#5300-05). FITC-conjugated streptavidin was purchased from BD Biosciences (Cat#554060). Human COVID-19 convalescent serum samples were purchased from Ray Biotech (Atlanta, GA). HRP-conjugated donkey anti-human IgG antibody was obtained from Jackson Immunoresearch (Cat#709-036-098).

Biotinylated human ACE-2 was purchased from ACRO Biosystems (Cat#AC2-H82F9). Purified Spike S1 RBD protein from Ray Biotech (Cat#230-30162), Spike S1 (Cat#40591-V08H) and South African variant (Cat#40591-V08H10) proteins were purchased from Sino Biological.

### Mice

BALB/c mice (Taconic Biosciences Inc.) 2-3 months age (female) were purchased and housed in the Emory University Division of Animal Resources (DAR) facility and used according to the University IACUC guidelines.

### Construction and expression of GPI-anchored SARS-CoV-2 S1 RBD-GM-CSF fusion protein

We have constructed a fusion protein gene by joining the DNA sequence of S1 RBD (amino acids 319-541), mouse GM-CSF, and the GPI-anchor signal sequence from human CD59 as in Figure 1a. This construct was cloned into the pCHO 1.0 vector using the AvrII and BstZ17l sites (Invitrogen). The DNA construct was then transfected into CHO-S cells (Invitrogen) and selected with puromycin and methotrexate. Expression of both the S1 RBD and GM-CSF on the surface of transfected CHO-S cells was confirmed by flow cytometry using fluorophore conjugated antibodies against RBD, Clone MM57 (Sino Biologicals) and GM-CSF, Clone MP1-22E9 (BioLegend). To confirm whether the fusion protein binds to its cognate receptor ACE2, flow cytometry analysis using biotinylated human ACE2 (ACRO Biosystems) was used and verification that the RBD-GM-CSF fusion protein retains its ability to bind to its cognate receptor was confirmed. CHO-S cell clones transfected with the fusion protein were grown in large quantities using a 5L bioreactor and used for purification by mAb-affinity chromatography.

### PIPLC treatment of CHO-S cells

To test the ability of phosphatidylinositol-specific phospholipase C (PIPLC) to cleave the GPI-RBD-GM-CSF fusion protein from the CHO-S cells, 0.25× 10^6^ cells were treated with 0.2 U PIPLC (Sigma cat# 554406) in 250 μL HBSS containing 0.1% BSA for 2 hours at 37°C. After treatment, cells were washed twice with FACS buffer before staining using PE anti-mouse GM-CSF antibody (BD cat# 554406) or PE Rat IgG2a,κ isotype control (BioLegend cat# 400508).

### Purification of GPI-RBD-GM-CSF and GPI-IL-12

Frozen CHO-S cell transfectants stably expressing GPI-IL-12 (42) or GPI-RBD-GM-CSF fusion protein were lysed for 1 hr at 4 °C in lysis buffer (50 mM Tris, 20 mM Iodoacetamide, 5 mM EDTA, 0.2% Tween 20, 2 mM PMSF and protease inhibitor cocktail, pH 8.0) and membranes containing the GPI-proteins were collected by centrifugation at 17,000x g for 1 hr at 4°C. The membranes were lysed with octyl glucoside (36) and GPI-RBD-GM-CSF and GPI-IL-12 present in the lysates were purified using anti-mouse GM-CSF antibody (Clone MP1-22E9, BioXCell) and anti-mouse IL-12 antibody (clone C17.8) coupled to NHS-Sepharose affinity (Cytiva) columns, respectively.

### SDS-PAGE and Western blot

Proteins were separated on a NuPAGE™ 12% Bis-Tris Gel, 1.0 mm x 10 well (Cat# NP0341BOX), and stained with Invitrogen’s Colloidal Blue Staining Kit, “Stain NuPAGE, Novex Bis-Tris Gel” (catalog # 46-7015, 46-7016), following manufacturer’s instructions. For Western blotting, proteins separated by 12% SDS Polyacrylamide gel electrophoresis (SDS-PAGE) under non-reducing condition are transferred onto a nitrocellulose membrane using semi-dry transfer apparatus via an electrical current. After transfer, the membranes with the proteins were blocked for 1 hr at room temperature with 5 % milk in phosphate buffered saline and 0.2 % Tween 20 (PBS-T) and incubated with a primary antibody (anti-RBD, anti-mouse GM-CSF or anti-mouse IL-12) overnight at 4 °C in PBS-T with on a shaker at low speed. Next day, membranes were washed three times with PBS-T and then incubated with appropriate secondary antibody conjugated with alkaline phosphatase that provides a visual color change upon addition of the chromogenic substrate (mixture of BCIP (5-bromo-4chloro-3-indolyl phosphate-catalog# 34040) and NBT (nitro-blue tetrazolium chloride, catalog# 34035 from *Thermo Scientific*)).

### Hybrid vaccine preparation by protein transfer of GPI-RBD-GM-CSF fusion protein and GPI-IL-12 onto VLP

Influenza VLPs containing codon-optimized hemagglutinin (H1 HA) and matrix M1 proteins derived from A/Puerto Rico/8/1934 (PR8) were produced in insect cells (Sf9) as described (40) and purified by tangential flow diafiltration and anion exchange (Capto Q) chromatography by Medigen (Frederick, MD). We incorporated the purified GPI-RBD-GM-CSF fusion protein along with GPI-IL-12 into PR8 influenza VLPs by protein transfer to prepare our VLP-RBD-GM-CSF-IL-12 vaccine. Protein transfer was performed by incubating 1 mg of enveloped influenza VLPs with 200 μg of purified GPI-RBD-GM-CSF and 25 μg of purified GPI-IL-12 at 37°C for 1 hour. Unincorporated GPI-cytokines were washed out by ultracentrifugation at 210,000 x *g* at 4°C and the resulting pellet was resuspended in DPBS. Resulting incorporation was detected by western blot and flow cytometry analysis using anti-RBD mAb, clone MM57 (Sino Biologicals), anti-mouse GM-CSF (clone MP1-22E9, BioLegend), and anti-mouse IL-12 (clone C17.8, Invitrogen) antibodies.

### Bone marrow derived dendritic cell stimulation assay

BMDCs were generated according to established protocols (43). Briefly, femurs of female BALB/c mice were removed and cleaned from surrounding muscle tissue. Bone marrow was flushed using RPMI-1640 medium with a 22G needle and syringe. Red blood cells (RBC) were lysed using RBC lysis buffer (MilliporeSigma, Burlington, MA, USA) and resulting cells were cultured in complete RPMI-1640 medium containing various concentrations of recombinant murine GM-CSF (rGM-CSF, BioLegend, San Diego, CA, USA), GPI-GM-CSF, GPI-RBD-GM-CSF or VLPs incorporated with GPI-RBD-GM-CSF at a density of 2 × 10^5^ cells/mL. After 3 days of culture, XTT assay reagent added, and absorbance measured after 3 hours at 480 nm and 660 nm for background reference.

### Immunization with VLP vaccine

BALB/c mice were administered with VLP vaccine, control VLP or PBS subcutaneously (100 μl volume per mouse) or intramuscularly (50 μl per mouse in the hind leg). Booster dose was given after 2 or 4 weeks. VLP vaccines were diluted in sterile PBS before administering to the mice.

### Enzyme-linked immunosorbent assay (ELISA)

SARS-CoV-2 S protein RBD specific antibodies of different subtypes (IgG, IgG1, IgG2a) were determined in sera by enzyme-linked immunosorbent assay (ELISA) using an approach similar to one previously described using PR8 or WSN as targets (44, 45). Briefly, 96-well ELISA plates were coated with 3 μg/ml GPI-RBD-GM-CSF in coating buffer (BioLegend) overnight at 4 °C. In some experiments, Spike S1 protein from commercial sources (RayBiotech or Sino Biologicals) was used as indicated in the figure legends. Plates were washed using a plate washer and washing buffer (PBS with 0.05% Tween20). Plates were blocked with assay diluent (PBS with 3% BSA) for 2 hours at room temperature on a rocker. Plates were washed again as above and diluted serum samples were added and incubated for another 2 hours at room temperature. Plates were washed and diluted secondary antibody (HRP-conjugated) against mouse total IgG or immunoglobulin isotypes (IgG1, IgG2a) were added and incubated for 30 minutes at room temperature. The plates were washed and TMB substrate solution (Cat# 555214, BD Biosciences) was added to the wells for color development. Reaction was stopped by adding 2N H_2_SO_4_ and read the absorbance at 450 nm.

To measure influenza antigen-specific antibody levels in immune sera, inactivated A/PR8 H1N1 virus (200 ng/well) was coated onto ELISA plates, followed by addition of diluted immune sera. IgG isotypes were measured using goat anti-mouse immunoglobulin (Ig) G, IgG1 and IgG2a, and horse-radish peroxidase (HRP)-conjugated secondary antibodies (Southern Biotechnology). Color reactions were developed with tetramethylbenzidine substrates (TMB, Invitrogen). Antibody levels are presented as optical density absorbance values at 450 nm (BioTek ELISA plate reader).

### Flow cytometry analysis of antibody response

For detection of Spike RBD-GM-CSF fusion protein on CHO-S cells, CHO-S cells (2-3×10^6^/ml) were incubated with anti-mouse GM-CSF antibody, anti-spike RBD (clone MM57, Sino Biological) in FACS buffer (PBS containing 2% BCS, 5 mM EDTA and 0.05% sodium azide), for 30-60 minutes on ice. For mouse sera, CHO-S cells were incubated with diluted serum samples (100-100,000 times diluted in FACS buffer). FITC-conjugated goat anti-mouse IgG/IgM was added as secondary antibody after washing off the unbound primary antibody. After incubation with secondary antibody, cells were washed with FACS buffer and resuspended in FACS buffer and acquired in a FACSCalibur (BD Biosciences) flow cytometer. Data analyzed using FlowJo software (FlowJo LLC).

### Focus Reduction Neutralization Assay

Live-virus SARS-CoV-2 neutralization antibodies were assessed using a full-length mNeonGreen SARS-CoV-2 (2019-nCoV/USA_WA1/2020), generated as previously described (46). FRNT-mNG assays were performed as previously described (39). Briefly, samples were diluted at 3-fold in 8 serial dilutions using DMEM (VWR, #45000-304) in duplicates with an initial dilution of 1:10 in a total volume of 60 μl. Serially diluted samples were incubated with an equal volume of SARS-CoV-2-mNG (100-200 foci per well) at 37° C for 1 hour in a round-bottomed 96-well culture plate. The antibody-virus mixture was then added to Vero cells and incubated at 37° C for 1 hour. Post-incubation, the antibody-virus mixture was removed and 100 μl of prewarmed 0.85% methylcellulose (Sigma-Aldrich, #M0512-250G) overlay was added to each well. Plates were incubated at 37° C for 24 hours. After 24 hours, methylcellulose overlay was removed, and cells were washed three times with PBS. Cells were then fixed with 2% paraformaldehyde in PBS (Electron Microscopy Sciences) for 30 minutes. Following fixation, plates were washed twice with PBS and foci were visualized on a fluorescence ELISPOT reader (CTL ImmunoSpot S6 Universal Analyzer) and counted using Viridot (47). The neutralization titers were calculated as follows: 1 -(ratio of the mean number of foci in the presence of sera and foci at the highest dilution of respective sera sample). Each specimen was tested in duplicate. The FRNT-mNG_50_ titers were interpolated using a 4-parameter nonlinear regression in GraphPad Prism 8.4.3.

### Plaque Reduction Neutralization Test (PRNT)

The titers of anti-SARS-CoV-2 neutralizing antibodies were measured in the serum of BALB/c mice using PRNT assay as described previously (41). Serum was diluted serially from 1:4 to 1:1024 and PRNT was conducted by using Wuhan strain of SARS-CoV-2 (BEI NR-52281). The highest dilution of serum resulting in 50% reduction in the number of plaques compared to the growth of the virus control was determined.

### Hemagglutination inhibition (HAI) assay

To assess the ability of immune sera to inhibit HA activity, we performed HAI assay using immune sera. The immune sera from each group were treated with receptor destroying enzymes (RDE, Sigma-Aldrich) for 18 h at 37 °C. Sera were incubated at 56 °C for 30 min for inactivation of Complement, followed by 10-fold serial dilutions in PBS. The serially diluted sera were incubated with 4 HA units of A/PR8 H1N1 virus for 30 min at room temperature, and then 0.5% chicken red blood cells (Lampire Biological Laboratories) were added to determine HAI titers. HAI titers were determined as the highest dilution factor inhibiting the formation of buttons with 0.5% chicken red blood cells.

### SARS-CoV-2 virus challenge

Female BALB/c mice (10-week-old) were purchased from Taconic Biosciences and housed in Emory University DAR facility. Mice were immunized with the VLP or VLP vaccine with cytokine adjuvants containing Spike RBD and given a booster dose 33 days after the first dose (n=10 mice per treatment). Blood was collected 2 weeks after the booster dose for antibody titer. Mice were transferred 3 months after the booster dose to ABSL-2 (n=5 for each treatment) for challenge with influenza A virus or ABSL-3 (n=5 for each treatment) for challenge with mouse-adapted SARS-CoV-2 virus at the Georgia State University (GSU). All the animal infection experiments using SARS-CoV-2 virus were conducted in a certified Animal Biosafety Level 3 (ABSL-3) laboratory at GSU. The protocol was approved by the GSU Institutional Animal Care and Use Committee (Protocol number A20044). Mice were inoculated intranasally with 10^5^ plaque-forming units (PFU) of mouse-adapted SARS-CoV-2 MA10 (48, 49). Animals were weighed and their appetite, activity, breathing, and neurological signs assessed twice daily. On day 3 after infection, animals were anesthetized with isoflurane and perfused with cold PBS. Lungs were collected, and flash-frozen in 2-methylbutane (Sigma, St. Louis, MO, USA).

### Quantification of the SARS-CoV-2 virus load in the lungs

Quantitative RT-PCR was used to measure viral RNA levels with primers and probes specific for the SARS-CoV-2 N gene as described previously (41). Viral genome copies were determined by comparison to a standard curve generated using a known amount of RNA extracted from previously titrated SARS-CoV-2 samples. Frozen tissues harvested from mock and infected animals were weighed and lysed in RLT buffer (Qiagen), and RNA was extracted using a Qiagen RNeasy Mini kit (Qiagen, Germantown, MD, USA). Total RNA extracted from the tissues was quantified and normalized, and viral RNA levels per μg of total RNA were calculated.

### Influenza A/PR8 H1N1 virus challenge

Three months after booster dose, BALB/c mice were challenged with a lethal dose of influenza A/PR8 H1N1 virus (10× LD_50_). After challenge, the mice were monitored for 14 days to record body weight changes and survival rates. To determine the protective efficacy and T cell responses after A/PR8 H1N1 infection, an additional set of immunized BALB/c mice was euthanized at day 5 post infection and lung tissues were collected for further analysis.

### Influenza virus titration in the lung

The lungs of the immunized BALB/c mice were harvested at day 5 after A/PR8 H1N1 infection and ground mechanically in 1.5 ml of PBS per lung. The lung extracts and lung cells were separated after centrifugation. Embryonated chicken eggs (Hy-Line North America, LLC) were incubated at 37 °C for 9–12 days to be inoculated with 10-fold serially diluted lung extracts. The virus titers were determined by hemagglutination assay of the allantoic fluids collected after 3 days of incubation. Virus titers as 50% egg infection dose (EID50)/ml were evaluated according to the Reed and Muench method (50).

### Statistical analysis

GraphPad Prism (version 9.1.0) was used to generate the graphs and analysis of the statistical significance as indicated in the figure legends. P value <0.05 is considered significant.

## Author contributions

Conceptualization: PS, CP, RB, SR, MK, SMK, MS

Methodology: CP, SR, PS, RB, MK, SMK, LJ, SN, ANB, TVG, CNW, LL

Investigation: RB, LEM, JO, KHK, NB, SS, PK, JLB

Funding acquisition: PS, SJCR, CP, SMK

Project administration: CP, SR, PS, SMK, MK, MS

Supervision: RB, SMK, PS, SR, CP, MS, MK,

Writing – original draft: RB, SMK, MK, CP, SR

Writing – review & editing: PS, SMK, MK, MS

## Data and materials availability

All data are available in the main text or the supplementary materials.

