## Supplementary data for "Influenza virus-like particle-based hybrid vaccine containing RBD induces immunity against influenza and SARS-CoV-2 viruses"

### Supplementary Materials & Methods

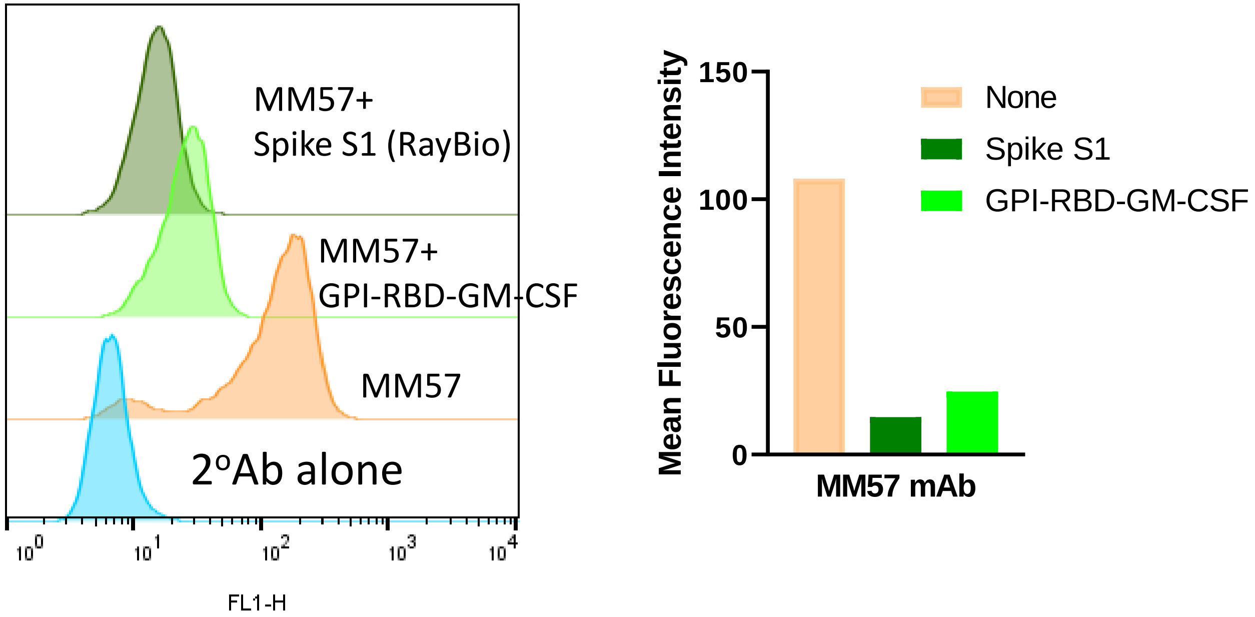


**Fig. S1. Purified GPI-RBD-GM-CSF fusion protein blocks binding of MM57 mAb to CHO-S cells expressing GPI-RBD-GM-CSF.** MM57 mAb (0.25 µg/ml) mixed with 3 µg/ml Spike S1 RBD (Ray Biotech; dark green) or GPI-RBD-GM-CSF fusion protein (light green bar) for 1 hr at room temperature in FACS buffer and then added to CHO-S cells and incubated for 1 hr at 4ºC. Washed the cells with FACS buffer 2 times and then incubated with FITC-conjugated secondary antibody for flow cytometry analysis.

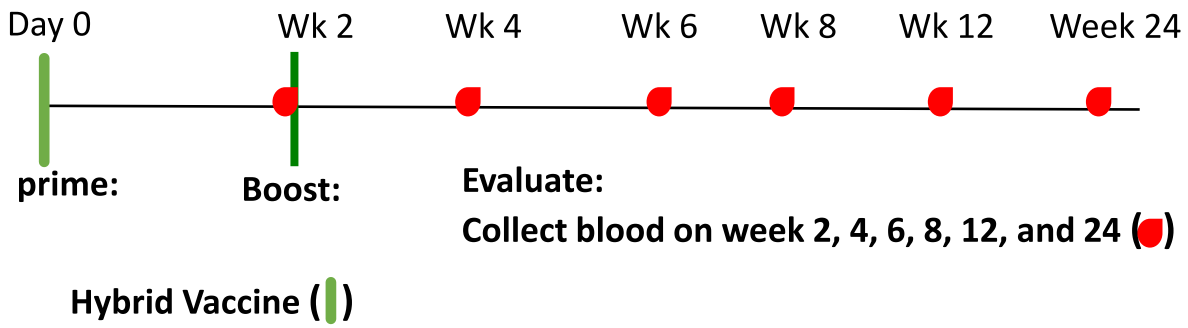


**Fig. S2. Design for VLP vaccination in BALB/c mice.** BALB/c mice (2-3 months old) administered with hybrid vaccine in PBS via subcutaneous (study 1) or intramuscular (study 2) route. Control mice received either PBS or influenza VLP. Purified GPI-RBD-GM-CSF or Spike RBD also administered for some groups. Booster dose was given 2 weeks after the first dose.

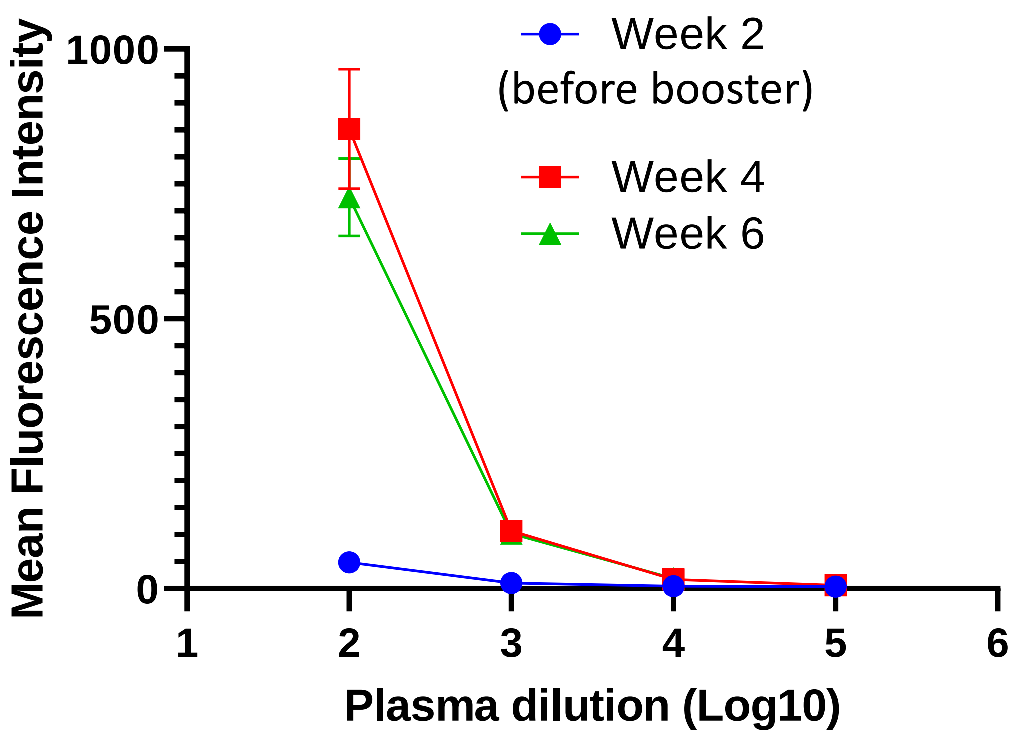


**Fig. S3. Booster dose induces high titers of anti-RBD antibody induction.** Mice were administered with hybrid vaccine (10 µg/dose) containing GPI-RBD-GM-CSF and GPI-IL-12 subcutaneously. Blood was collected after 2 weeks, and the following day booster dose was given. Blood was collected 2 weeks and 4 weeks after the booster dose (4 weeks and 6 weeks after the first dose). CHO-S cell transfectants expressing GPI-RBD-GM-CSF were used for detection of antibody by flow cytometry. FITC-conjugated anti-mouse IgM/IgG secondary antibody was used.

VLP vaccine 10 µg

Week 4, 100x dilution

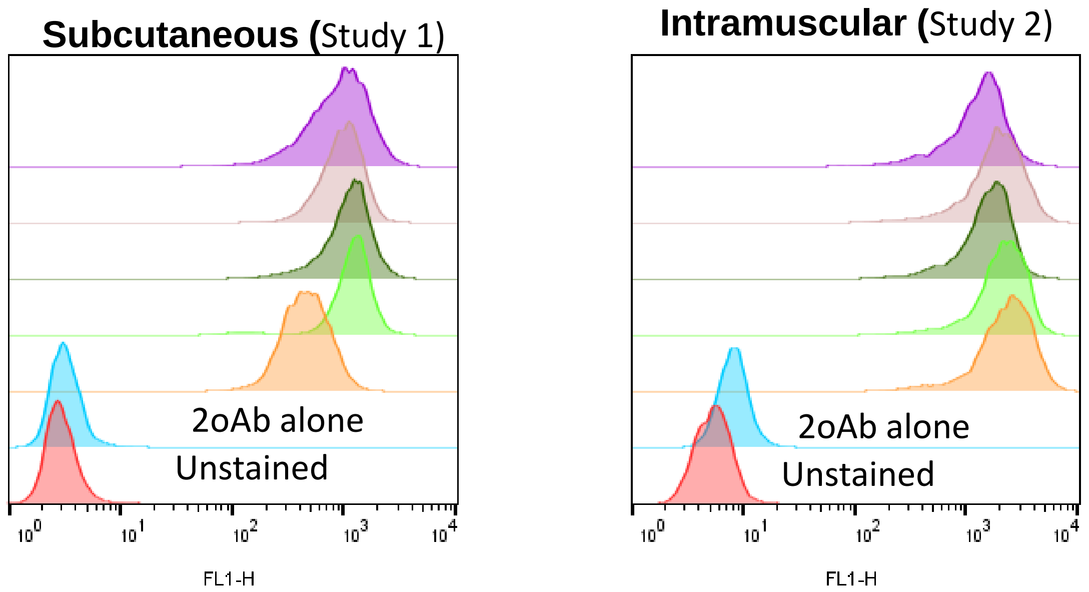


**Fig. S4. VLP vaccine induces comparable levels of antibody against RBD.** Mice (n=5) were administered with 10 µg hybrid vaccine containing GPI-RBD-GM-CSF and GPI-IL-12 either subcutaneous (left) or intramuscular (right) route. Booster dose was given 2 weeks later, and blood was collected 2 weeks after the booster dose (4 weeks after the first dose). CHO-S cell transfectants expressing GPI-RBD-GM-CSF were used for detection of serum antibody (1:100 dilution) by flow cytometry. FITC-conjugated anti-mouse IgM/IgG secondary antibody was used.

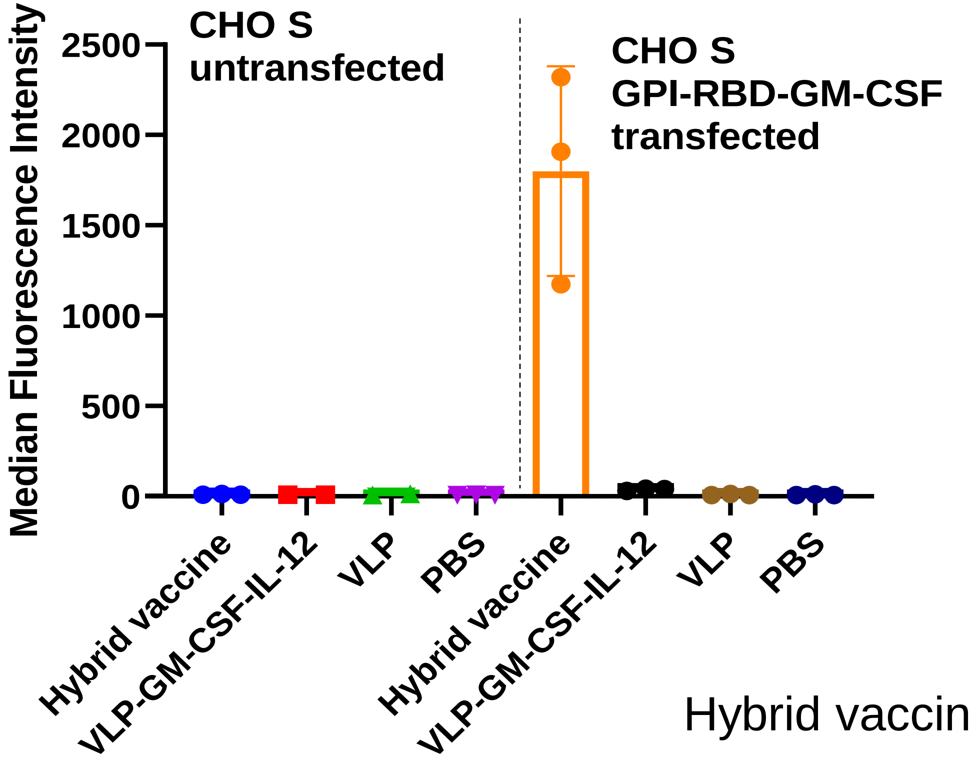


Hybrid vaccine: VLP-GPI-RBD-GM-CSF-IL-12

**Fig. S5**. **Hybrid vaccine induces S1-RBD specific antibody response.** Mice were administered with 10 µg VLP vaccine containing GPI-IL-12 and either GPI-GM-CSF or GPI-RBD-GM-CSF subcutaneous route. Booster dose was given 2 weeks later, and blood was collected 2 weeks after the booster dose (4 weeks after the first dose). CHO S cells (left) or CHO S transfectants expressing GPI-RBD-GM-CSF (right) were used for detection of serum antibody (1:50 dilution) by flow cytometry. FITC-conjugated anti-mouse IgM/IgG secondary antibody was used.

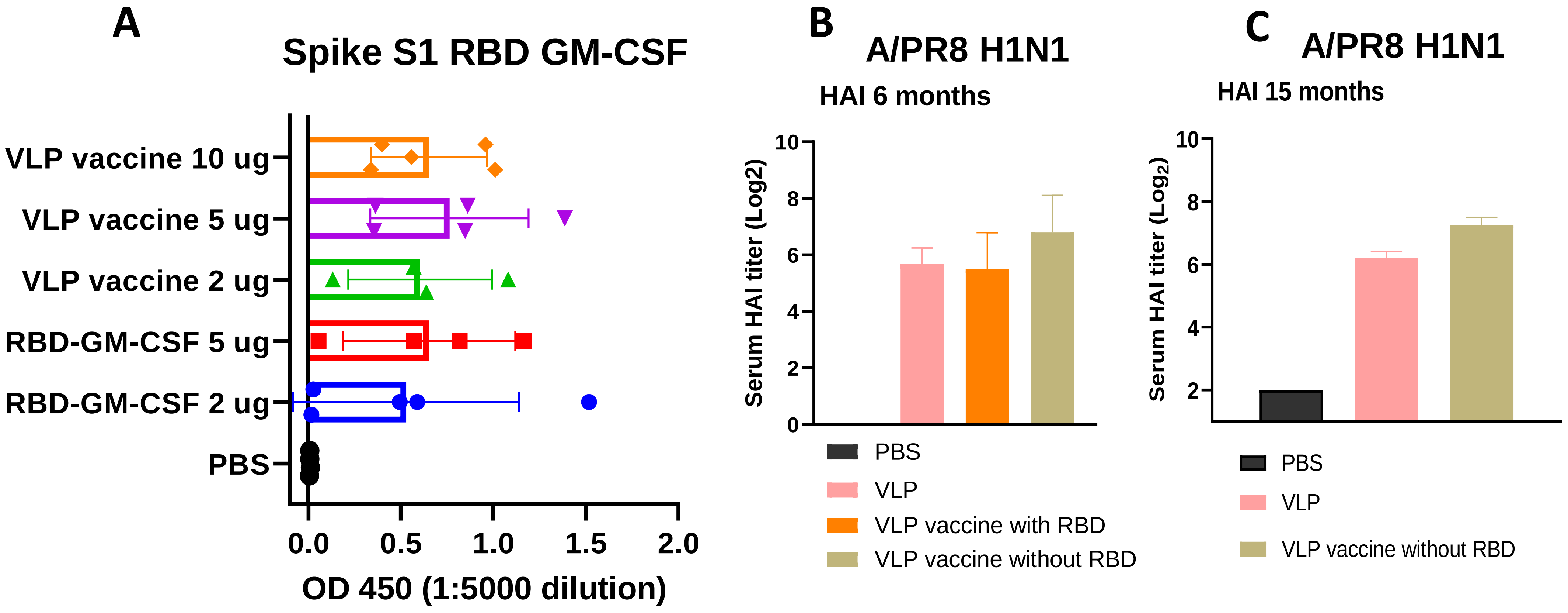


**Fig. S6: Antibody response maintained even after 6 months.** (A) ELISA plates were coated with GPI-RBD-GM-CSF. Serum samples from various groups of mice immunized with GPI-RBD-GM-CSF or VLP vaccine containing the GPI-RBD-GM-CSF and GPI-IL-12 were diluted 5000 times and added to the wells after blocking the plates. Goat anti-mouse IgG-HRP conjugate was used to detect the antibody levels. Hemagglutination inhibition (HAI) titer in the sera after 6 months (B) and 15 months (C) of vaccination.

.

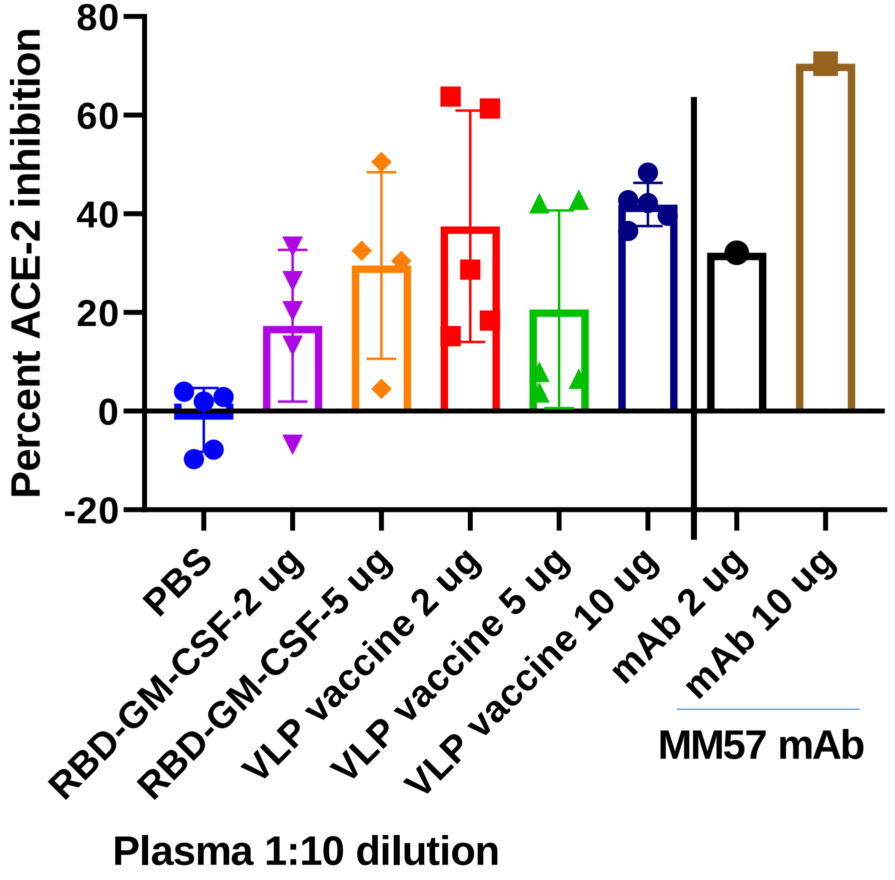


**Fig. S7**. **VLP vaccine-induced antibody blocks ACE-2 binding to RBD.** Mice administered with GPI-RBD-GM-CSF protein or hybrid vaccine subcutaneous route. Booster dose given 2 weeks later, and blood collected 6 weeks after the booster dose (8 weeks after the first dose). CHO S cell transfectants were incubated with diluted plasma or MM57 mAb followed by biotinylated ACE-2 (3 µg/ml). FITC-conjugated streptavidin (1:50 dilution) used for detecting the cell surface RBD-bound ACE-2.
Hybrid vaccine: VLP incorporated with GPI-RBD-GM-CSF and GPI-IL-12

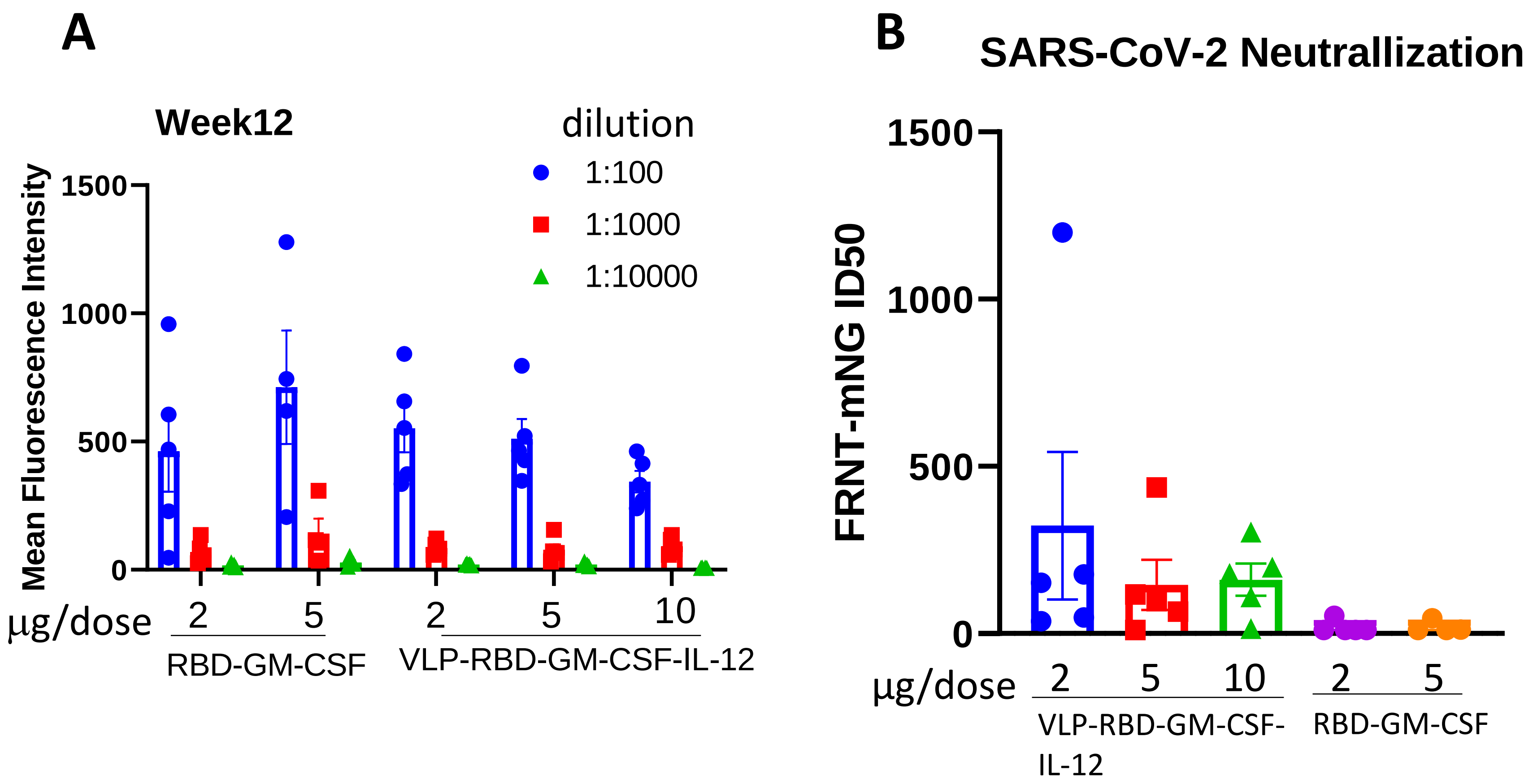


**Fig. S8. Hybrid vaccine induces virus-neutralizing antibody.** (A) Antibody levels in the serum detected by flow cytometry using CHO-S cells as described in Methods. (B) Microneutralization of SARS-CoV-2 infection using Vero6 cells performed by pre-incubating the WA11 strain of SARS-CoV2 with serially diluted heat-inactivated plasma samples. .

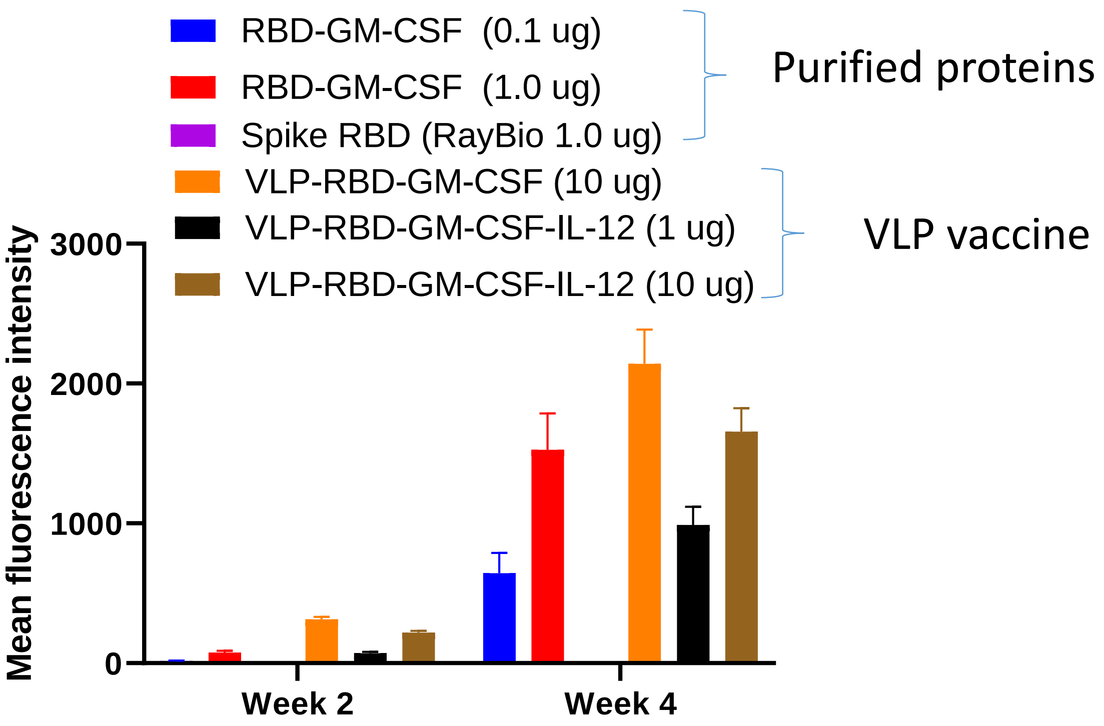


**Fig. S9. GM-CSF and VLP enhance antibody response against RBD.** Mice administered with RBD-GM-CSF, VLP vaccine containing GPI-RBD-GM-CSF with or without GPI-IL-12 intramuscularly. Blood was collected after 2 weeks (week 2), and the following day booster dose was given. Blood collected 2 weeks after the booster dose (week 4: 4 weeks after the first dose). CHO S cell transfectants expressing GPI-RBD-GM-CSF used for detection of anti-RBD antibody by flow cytometry. FITC-conjugated anti-mouse IgM/IgG secondary antibody used. Spike RBD from Ray Biotech (1.0 µg) was used as a control protein antigen. Sera dilution is 100x.

Note: No detectable antibody in mice administered with Spike RBD from RayBiotech

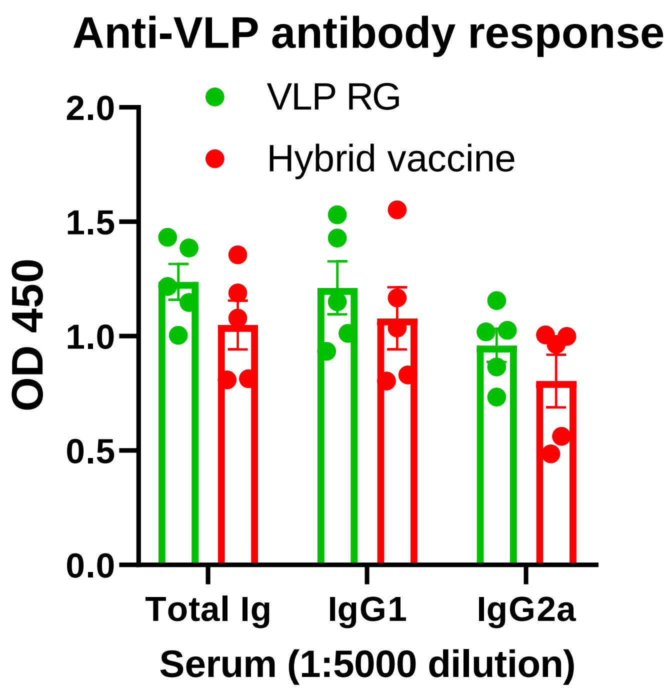


**Fig. S10**. **Hybrid vaccine induces antibody response against influenza VLP antigens.** ELISA plates coated with influenza VLP. Serum samples (12 weeks after first dose) from mice immunized with VLP RG (hybrid vaccine without GPI-IL-12) or hybrid vaccine were diluted 5000 times and added to the wells after blocking the plates. Total IgG and isotype specific anti-mouse IgG-HRP conjugate used to detect the bound antibody.
Hybrid vaccine: VLP incorporated with GPI-RBD-GM-CSF and GPI-IL-12

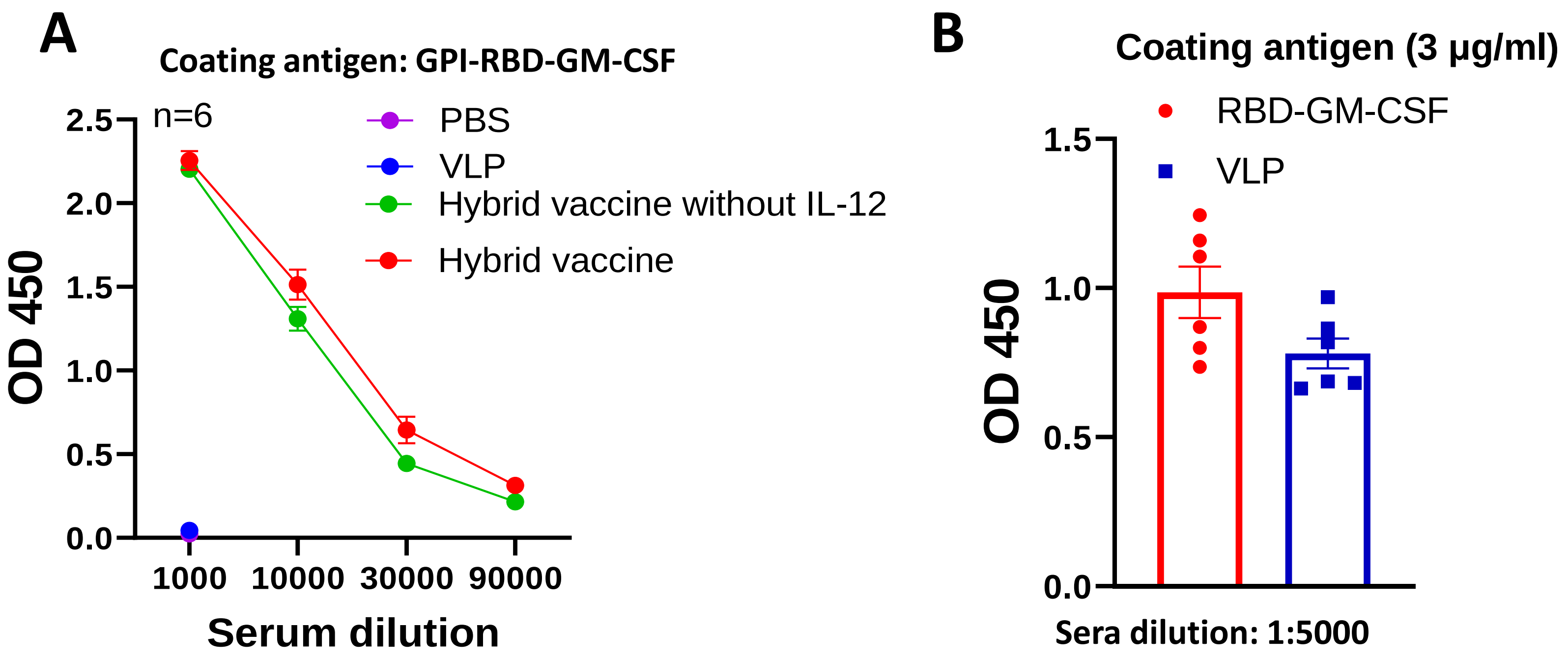


**Fig. S11**. **Hybrid vaccine induces antibody response against RBD-GM-CSF and influenza VLP antigens.** ELISA plates coated with GPI-RBD-GM-CSF (A & B) or influenza VLP (B). Serum samples (2 weeks after the booster dose) from mice immunized with PBS, VLP, hybrid vaccine or hybrid vaccine without IL-12 (A) or hybrid vaccine (B) diluted and added to the wells after blocking the plates. Anti-mouse IgG-HRP conjugate used to detect the bound IgG antibody
